# Inulin nanostructures: The sweet-spot of carbohydrate self-assembly

**DOI:** 10.1101/2021.07.13.451276

**Authors:** Zohar A. Arnon, Márkó Grabarics, Topaz Kreiser, Avi Raveh, Kevin Pagel, Ehud Gazit

## Abstract

Supramolecular architecture formation by the self-assembly of proteins and nucleic acids is well studied. Yet, the spontaneous organization of oligosaccharides, the most common polymers in nature, is less explored. Here, using inulin as a model, we identify the “sweet spot” length for oligosaccharide assembly. Inulin forms discrete spheres in a concentration-dependent manner. Size-based fractionation displayed markedly different aggregation morphologies. Based on these findings, we believe that carbohydrates could become an important source for novel self-assembling materials.

## Introduction

In the field of materials science, there is an increasing interest in the application of bioorganic materials. Specifically, molecular self-assembly is an attractive approach to produce novel materials. The design and fabrication of ordered supramolecular structures through molecular self-assembly are often inspired by elegant biological architectures. These natural systems offer a diverse set of attributes, frequently desired for nanotechnological applications. Traditionally, the two main avenues of bio-inspired self-assembly research have utilized polymers of amino acids (peptides and proteins) or nucleotides (DNA and RNA)^1–6^. However, the most abundant organic polymers on Earth are oligo- and polysaccharides, which are currently sparsely used as self-assembling materials, despite their diversity in size and physico-chemical properties. For instance, all plants use cellulose to build their cell walls^7^; amylum, better known as starch, is used by most green plants for energy storage^8^. In addition to their abundance in nature, there is a broad range of properties that can be attained by a different combination of polysaccharides. For example, polysaccharides are accountable for the soft properties of cotton, as well as for the rigid and tough bark of the giant sequoia trees, mostly due to different cellulose/lignin ratios^9,10^. Recently, the field of self-assembling saccharide-based materials is gaining increased attention^11,12^. Synthesized di- and hexasaccharides were shown to self-assemble into nanostructures of varying morphologies, which surprisingly display intrinsic fluorescence within the visible spectrum in an excitation-dependent manner^13^. Synthetic saccharides were also used as functional materials for applications such as biofilm inhibition, drug and supplement delivery and tissue engineering^14–16^. Yet, considering their abundance in nature, saccharides remain overlooked in materials science and their exceptional structural properties are still not fully exploited. Generally, for the spontaneous process of molecular self-assembly, the building blocks first must be dissolved to allow for reorganization into assemblies through non-covalent interactions with favorable energetic states^17^. However, if the molecule is too easily dissolved in the medium, the individual building blocks are relatively energetically stable, and very high concentrations would be required for reorganization and subsequent assembly. Typical examples are mono- and disaccharides such as glucose and sucrose, which are highly soluble in water^18^. In contrast, long polysaccharide chains such as cellulose, are insoluble in water due to the extensive network of hydrogen bonds between individual polymer chains. The intramolecular interactions between small mono- and disaccharide molecules are generally too weak to stabilize the structure, while the numerous interactions of polysaccharides such as cellulose are too strong to disassociate into monomers in aqueous solution. Therefore, we were looking for a polymer with intermediate solubility in water, and with an intermediate degree of polymerization. Hence, inulin, an unbranched oligosaccharide with degree of polymerization (DP) of 2-60, was investigated for its self-assembly propensity (Fig. 1). Here, we aim to identify the optimal range of the molecular solubility-size scale, where ordered supramolecular structures are formed, *i.e.* the sweet-spot of carbohydrate self-assembly.

**Fig. 1.**
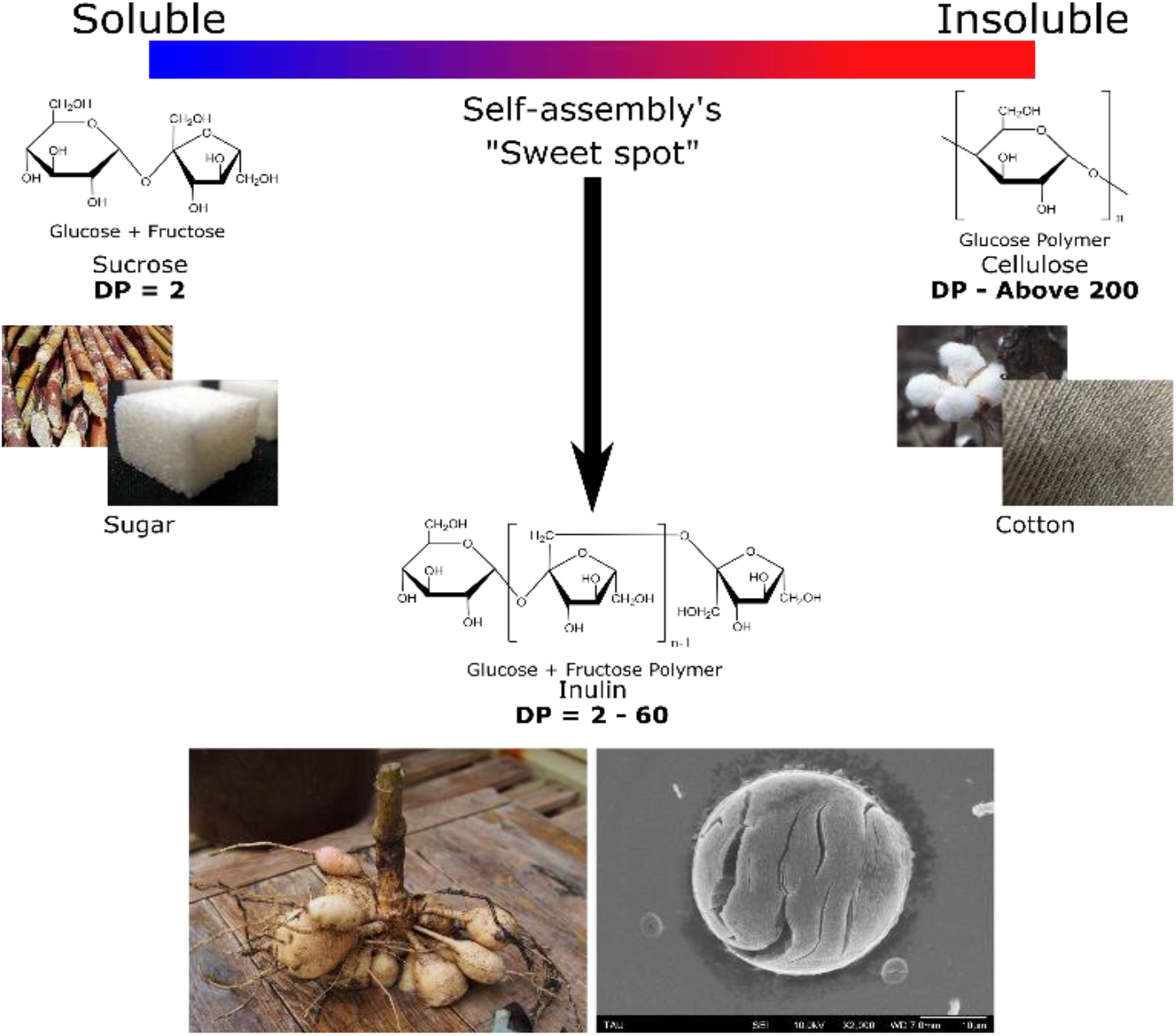
Solubility optimization of oligosaccharide self-assembly. The solubility of oligosaccharide polymers varies greatly with the degree of polymerization (DP). For example, sucrose, (DP = 2) found in sugarcane is highly soluble in water, while cellulose (DP of hundreds to thousands), commonly found in cotton, is poorly soluble. Inulin, extracted from dahlia tubers, is in between in the solubility scale, and could have the desired solubility for self-assembly.

## Results

Inulin powder, extracted from dahlia tubers, was dissolved in water at different concentrations. Due to the heterogeneous composition of the powder, the exact solubility cannot be determined unambiguously. Since the solubility of saccharides increases significantly with elevated temperature, water at 90 °C allows to easily dissolve inulin to a high concentration of 50 mg/ml, while at room temperature the solubility is approximately 5 mg/ml. Allowing a hot solution of highly concentrated inulin to cool down to room temperature therefore induces the self-association of molecules within solution as the solubility drops. The turbidity of the cooled down assembled solutions was evaluated by measuring the absorbance at 405 nm. As expected, in correlation with increasing concentration, the inulin assemblies lead to a more turbid solution (Fig. 2a). Moreover, the resulting assemblies were found to be denser at a higher initial concentration of the solution, which also affects the resulting morphology. Optical microscopy revealed that the assemblies blend and entangle to create rugged planes at higher concentrations, while low concentrations give rise to mostly discrete assemblies, with some entangled planes (Fig. 2b). Electron microscopy allowed a closer inspection of the assemblies (Fig. 2c). At lower concentrations, the assemblies are more dispersed, and discrete spheres are abundant throughout the sample. The self-assembled architecture is important, specifically shown previously for spheres, since the spherical architecture of self-assembled materials, in the case of peptide self-assembly, exhibited enhanced mechanical properties and remarkable optical scattering^19,20^.

**Fig. 2.**
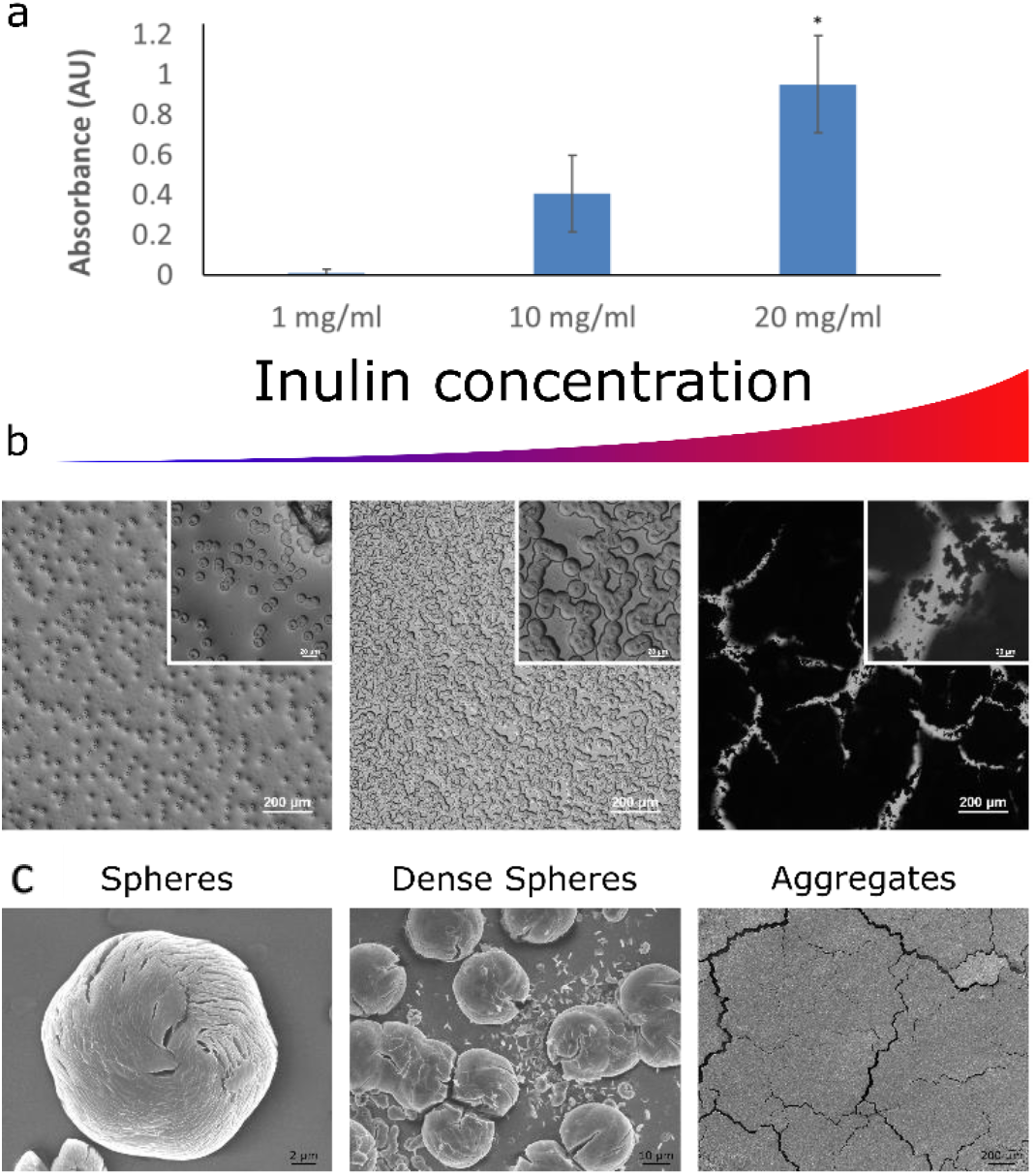
Inulin concentration dependent assemblies. (a) Structures formation was induced by dissolving inulin in water at 90 °C and cooling down the solutions gradually. The presence of the assemblies led to increased turbidity of the solution in correlation with the initial inulin concentration. As can be seen from optical (b) and electron (c) microscopy, lower concentrations of the oligosaccharide lead to the formation of discrete spherical structures, while higher concentrations lead to dense and entangled arrays of indistinct assemblies.

We then aimed to achieve a more homogenous sample by fractionating a solution of the dissolved oligosaccharide. Ultrafiltration membranes with a cut-off filter of 10 kDa were used, and the fractions below and above the cut-off value, including an untreated sample as a control were investigated. Nano-electrospray ionization mass spectrometry (nESI-MS) experiments were performed in negative ion mode to evaluate the ultrafiltration process. This technique enables the determination of the mass of molecular ions generated from a solution. Assuming that the relative intensities of ions in the mass spectra are representative of the relative concentrations of their parent molecules in solution, the size and molecular weight distribution of the samples can be reconstructed (Fig. 3a and 3b). Our results show a clear difference between the molecular size distribution of the two membrane-filtered fractions and the unfiltered inulin sample. The number average molecular mass (*M*_*n*_) and weight average molecular mass (*M*_*w*_) of the fraction below the cut-off limit appear as 3000 Da and 3221 Da, respectively, while the average degree of polymerization *(DP*_*avg*_) is 18.4. In comparison, the respective values of the fraction above cut-off are *M*_*n*_ = 4255, *M*_*w*_ = 4629 and *DP*_*avg*_ = 26.1. We believe that the above cut-off sample exhibits average molecular masses lower than 10 kDa (the supposed cut-off) due to the linear nature of the polymer chains – as far as possible from ideal globular particles. In accordance with expectations, the representative measures of the unfiltered inulin sample fall between those of the two membrane-filtered fractions: *M*_*n*_ = 3566, *M*_*w*_ = 3917 and *DP*_*avg*_ = 21.9 (see Table S1). Importantly, the differences in the molecular weight distribution of the various fractions lead to assemblies with markedly distinct morphologies (Fig. 3c).

**Fig. 3.**
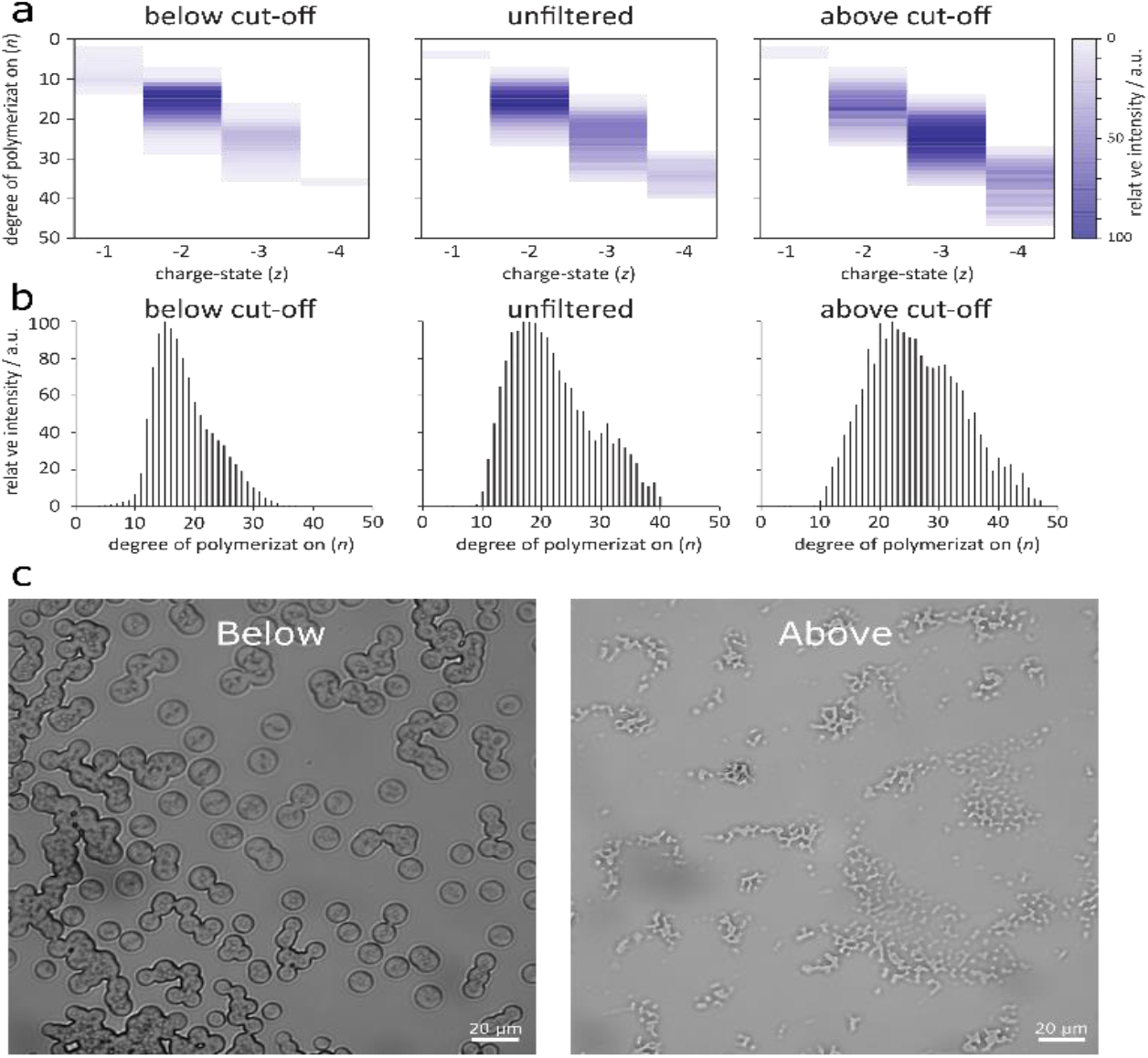
Membrane-filtered inulin fractions – size distribution and morphology. (a) Matrices depicting the relative intensities of inulin ions as a function of polymerization degree and charge state in nESI-MS experiments. (b) Molecular size distribution of various inulin fractions reconstructed from MS measurements. (c) Optical microscopy images of the spherical structures formed from the shorter inulin polymer chains (below), and the amorphous aggregates of the longer polymer chains (above). Both samples at a concentration of 1 mg/ml.

Additional information on MS data evaluation and statistical characteristics (including definitions) of the reconstructed distributions can be found in the Supplementary Material.

In order to ensure that both the above and below cutoff samples are holding the same concentration of polymer, we lyophilized the samples to achieve post-fractionation dry powders. These powders were then re-dissolved individually in the desired concentrations. Since we considered weight concentrations (mg/ml), the number of polymer chains is different between samples, but the amount of monosaccharide units is similar in both solutions. In this case, the differences between the below and above fractions were visible at lower concentrations of the samples, as at high concentrations the resulting assemblies were too dense. At a concentration of 1 mg/ml, the shorter polymeric chains assembled into relatively large and discrete spherical morphologies. Contrarily, the association of the longer polymers resulted in amorphous clusters of matter (Fig. 3c). This indicates that the inulin polysaccharides with a DP of ~ 20 are favorable for self-association in aqueous solution.

## Conclusions

To conclude, as in nature, saccharides have the potential to serve as a significant class of self-assembled materials in the fields of nanotechnology and materials science. To fully realize their potential, an in-depth understanding and control over the assembly process is essential. Here, we investigated the molecular details of the assembly process and in particular the effect of polymer length. Our results show that there is indeed a sweet spot for the self-assembly of oligosaccharides. We clearly demonstrate that smaller inulin size fractions form well-defined supramolecular assemblies, while large fractions assemble into amorphous aggregates. We hope this work will serve as a conceptual stepping-stone towards establishing polysaccharides as an important class of self-assembling materials for nanotechnological applications.

## Experimental Section

### Turbidity assay

Inulin (Sigma-Aldrich-I3754) in various concentrations was dissolved in ddw and heated to 90 °C until fully dissolved. 200 μL of the heated solutions were added to a 96-well plate in quartets. A hot ddw-only solution served as a control. The plate was sealed (TempPlate^®^ sealing film, USA scientific) and kept in room temperature for 3 weeks. Then, turbidity absorbance was measured at 405 nm in CLARIOstar plate reader (BMG).

### High-resolution Scanning Electron Microscopy

Inulin structures formation was induced by dissolving inulin in double distilled water (ddw) at 90 °C and cooling down the solutions gradually. Subsequently, 10 μL samples of these solutions were placed on a glass slide and left to dry at room temperature. Samples were then coated with chrome and viewed using a JSM-6700 field-emission HR-SEM (Jeol, Tokyo, Japan), equipped with a cold field emission gun, operating at 10 kV.

### Structures ultrafiltration

Ultrafiltration membranes (Merck) with a cut-off filter of 10 kDa were used to fractionate the sample. A sample of 10 mg/ml of the dissolved polysaccharide is transferred through the membrane using a centrifuge and both the sample that is passed through the filter and the sample that does not pass are collected.

### Nano-electrospray ionization mass spectrometric (nESI-MS)

The experiments were performed in negative ion mode using a modified Synapt G2-S HDMS Q-ToF instrument (Waters Corporation, Manchester, UK), equipped with a radiofrequency-confining drift cell. Lyophylized solid inulin samples were dissolved in water/methanol 50:50 (*v:v*) with 0.1% formic acid at a concentration of 1 mg/mL. For each analysis, 5 μL of sample solution was loaded into Pt/Pd-coated borosilicate capillaries prepared in-house. Ions were generated using a nano-electrospray (nESI) ion source, applying 0.6-0.7 kV voltage to the capillaries. All measurements were performed in the range of *m/z* 100–5000. Typical instrument settings were as follows: capillary voltage 0.6-0.7 kV, source temperature 40 °C, sampling cone voltage 110 V, cone offset 70 V, trap collision energy 4 V, transfer collision energy 2 V, trap gas flow 2 mL/min. All solvents and reagents employed were of LC-MS grade.

## Supporting Information

Supporting Information is available online or from the author.

## Author contributions

Z.A.A., M.G., T.K., K.P., and E.G. conceived and designed the experiments. Z.A.A., M.G., T.K., performed the experiments. Z.A.A., M.G., T.K., A.R., K.P., and E.G. wrote the manuscript. M.G. performed the MS experiments and analysis. All authors discussed the results, provided intellectual input and critical feedback and commented on the manuscript.

## Acknowledgements

We thanks all our lab members for the fruitful discussions.

## References

1. E. Gazit, Nat. Chem., 2015, 7, 14–15.

2. M. Burnworth, L. Tang, J. R. Kumpfer, A. J. Duncan, F. L. Beyer, G. L. Fiore, S. J. Rowan and C. Weder, Nature, 2011, 472, 334–337.

3. Y. Krishnan and N. C. Seeman, Chem. Rev., 2019, 119, 6271–6272.

4. E. Krieg, M. M. C. Bastings, P. Besenius and B. Rybtchinski, Chem. Rev., 2016, 116, 2414–2477.

5. T. Aida, E. W. Meijer and S. I. Stupp, Science, 2012, 335, 813–817.

6. S. Zhang, Nat. Biotechnol., 2003, 21, 1171–1178.

7. R. J. Moon, A. Martini, J. Nairn, J. Simonsen and J. Youngblood, Chem. Soc. Rev., 2011.

8. H. F. Zobel, Starch - Stärke, 1988, 40, 44–50.

9. S. H. Lamlom and R. A. Savidge, Tree Physiol., 2006, 26, 459–468.

10. N. Abidi, L. Cabrales and C. H. Haigler, Carbohydr. Polym., 2014, 100, 9–16.

11. C. Gao and G. Chen, Acc. Chem. Res., 2020, 53, 740–751.

12. Z. Ma and X. X. Zhu, J. Mater. Chem. B, 2019, 7, 1361–1378.

13. Y. Yu, S. Gim, D. Kim, Z. A. Arnon, E. Gazit, P. H. Seeberger and M. Delbianco, J. Am. Chem. Soc., 2019, 141, 4833–4838.

14. G. Tao, T. Ji, N. Wang, G. Yang, X. Lei, W. Zheng, R. Liu, X. Xu, L. Yang, G. Q. Yin, X. Liao, X. Li, H. M. Ding, X. Ding, J. Xu, H. B. Yang and G. Chen, ACS Macro Lett., 2020, 9, 61–69.

15. G. A. Valencia, E. N. Zare, P. Makvandi and T. J. Gutiérrez, Compr. Rev. Food Sci. Food Saf., 2019, 18, 2009–2024.

16. S. Gim, Y. Zhu, P. H. Seeberger and M. Delbianco, Wiley Interdiscip. Rev. Nanomedicine Nanobiotechnology, , DOI:10.1002/wnan.1558.

17. T. O. Mason, T. C. T. Michaels, A. Levin, C. M. Dobson, E. Gazit, T. P. J. Knowles and A. K. Buell, J. Am. Chem. Soc., 2017, 139, 16134–16142.

18. L. A. Alves, J. B. Almeida E Silva and M. Giulietti, J. Chem. Eng. Data, 2007, 52, 2166–2170.

19. L. Adler-Abramovich, N. Kol, I. Yanai, D. Barlam, R. Z. Shneck, E. Gazit and I. Rousso, Angew. Chemie Int. Ed., 2010, 49, 9939–9942.

20. Z. A. Arnon, D. Pinotsi, M. Schmidt, S. Gilead, T. Guterman, A. Sadhanala, S. Ahmad, A. Levin, P. Walther, C. F. Kaminski, M. Fändrich, G. S. Kaminski Schierle, L. Adler-Abramovich, L. J. W. Shimon and E. Gazit, ACS Appl. Mater. Interfaces, 2018, 10, 20783–20789.

