## Supplementary material for "Inulin nanostructures: The sweet-spot of carbohydrate self-assembly": Table S1

### Supporting Information

#### Nano-electrospray ionization mass spectrometric (nESI-MS)

Data acquisition, processing, evaluation and visualization were performed using MassLynx v4.1 (Waters, Manchester, UK), OriginPro 2017 (OriginLab Corporation, Northampton, USA) and GlycoWorkBench[1] software. Briefly, mass peaks were assigned manually and the intensity of each peak was determined based on the peak area. The areas of monoisotopic peaks corresponding to the same ion were summed. For creating the matrices in Figure 3a, ions of different charge-states were treated separately. For Figure 3b, all ionic species corresponding to the same parent molecule in solution (same degree of polymerization) were added together – irrespective of the observed charge- state in the MS experiment – to represent the distribution of molecules in the samples. It is assumed that the relative intensities in the mass spectra are representative of the relative concentrations in solution, giving an accurate picture of the molecular weight distribution of the samples.

#### Molecular size distribution

Definitions of the statistical measures employed for the characterization of the molecular size distribution[2,3] of inulin are given below.

a) Number average molecular weight ($M_n$):

$$M_n = \frac{\sum_i M_i c_i}{\sum_i c_i}$$

Here, $M_i$ is the molecular weight of the $i$th component, while $c_i$ is its concentration. The actual concentrations can be substituted with the measured intensities in the mass spectra, assuming that they are proportional to the concentrations of the parent molecules in solution.

b) Weight average molecular weight ($M_w$):

$$M_w = \frac{\sum_i M_i {c_i}^2}{\sum_i M_i c_i}$$

c) Average degree of polymerization ($DP_{avg}$):

$$DP_{avg} = \frac{\sum_i DP_i c_i}{\sum_i c_i}$$

Here, $DP_i$ is the degree of polymerization of the $i$th component.

The characteristic measures of the size and molecular weight distribution of the various inulin samples are summarized below in Table S1.

**Table S1. Molecular size distribution of membrane-filtered and untreated inulin samples.**

| **Sample** | **Molecular weight** | | **Degree of polymerization** | |
|---|---|---|---|---|
| | **Number average** | **Weight average** | **Average** | **Mode** |
| Below cut-off | 3000 Da | 3221 Da | 18.4 | 15 |
| Unfiltered | 3566 Da | 3917 Da | 21.9 | 17 |
| Above cut-off | 4255 Da | 4629 Da | 26.1 | 22 |

#### Bibliography


1 A. Ceroni, K. Maass, H. Geyer, R. Geyer, A. Dell and S. M. Haslam, *J. Proteome Res.*, 2008, **7**, 1650–1659.

2 P. J. Flory, *J. Am. Chem. Soc.*, 1940, **62**, 1561–1565.

3 U. Bahr, A. Deppe, M. Karas, F. Hillenkamp and U. Giessmann, *Anal. Chem.*, 1992, **64**, 2866–2869.
